# Quantitative Isothermal Amplification on Paper Membranes using Amplification Nucleation Site Analysis

**DOI:** 10.1101/2022.01.11.475898

**Authors:** Benjamin P. Sullivan, Yu-Shan Chou, Andrew T. Bender, Coleman D. Martin, Zoe G. Kaputa, Hugh March, Minyung Song, Jonathan D. Posner

## Abstract

Quantitative nucleic acid amplification tests (qNAATs) are critical in treating infectious diseases, such as in HIV viral load monitoring or SARS-CoV-2 testing, in which viral load indicates viral suppression or infectivity. Quantitative PCR is the gold standard tool for qNAATs; however, there is a need to develop point-of-care (POC) qNAATs to manage infectious diseases in outpatient clinics, low- and middle-income countries, and the home. Isothermal amplification methods are an emerging tool for POC NAATs as an alternative to traditional PCR-based workflows. Previous works have focused on relating isothermal amplification bulk fluorescence signals to input copies of target nucleic acids for sample quantification with limited success. In this work, we show that recombinase polymerase amplification (RPA) reactions on paper membranes exhibit discrete fluorescent amplification nucleation sites. We demonstrate that the number of nucleation sites can be used to quantify HIV-1 DNA and RNA in less than 20 minutes. An image-analysis algorithm quantifies nucleation sites and determines the input nucleic acid copies in the range of 67-3,000 copies per reaction. We demonstrate a mobile phone-based system for image capture and onboard processing, illustrating that this method may be used at the point-of-care for qNAATs with minimal instrumentation.

## 1. Introduction

Nucleic acid amplification tests (NAATs) are critical tools in diagnosing infectious diseases due to their unparalleled specificity and sensitivity. Quantitative nucleic acid amplification tests (qNAATs) quantify target nucleic acids (NA) concentrations in biological samples. This is crucial in many applications, including HIV viral load testing where NA concentrations (i.e. viral load) are related to viral suppression and transmission risk. HIV-positive individuals with viral loads of less than 1,000 copies of viral RNA per mL of plasma have significantly better health outcomes and reduced risk of transmitting HIV to partners.^1–3^ Recently, SARS-CoV-2 viral load measurements have been related to infectivity, with higher viral loads being strongly correlated to cultivable virus.^4,5^ Quantification of pathogen load can be crucially important in triaging and determining appropriate clinical care.

Quantitative PCR (qPCR) is the gold standard for quantifying NAs and amplifies pathogenic nucleic acids using precise thermocycling for denaturation, annealing, and extension of duplicated DNA. qPCR provides quantification over 9 orders of magnitude relative to known quantities of DNA. Digital droplet PCR (ddPCR) has emerged as the most precise method for absolute NA quantification. It leverages Poisson statistics with thousands of discrete PCR amplification reactions to provide absolute sample quantification.^6,7^ ddPCR typically has lower dynamic range than qPCR (5+ orders of magnitude for commercial systems such as the BioRad QX200^8^) due to signal saturation and requires separate droplet generation prior to amplification cycling. Both qPCR and ddPCR are traditionally restricted to well-instrumented laboratories or hospitals due to cold chain dependent reagents, delicate instrumentation, reliable electrical power, proficient laboratory staff, and appropriate infrastructure to host required equipment.^9^

Pathogen quantification using qPCR or ddPCR in outpatient or low- and middle-income country clinics is challenging due to the logistics around specimen collection, transport, batched testing, and the return of results to clinicians and patients (in addition to the aforementioned restriction to centralized laboratories).^10^ In outpatient clinics, this can result in delayed diagnoses that prevent immediate linkage to appropriate treatment or lead to loss-to-follow-up.^11,12^ Consequently, there is an unmet need to develop inexpensive point-of-care (POC) qNAATs, particularly in low- and middle-income countries (LMIC).^9,10^ Isothermal amplification methods are emerging as an alternative to PCR, as they require significantly less instrumentation and produce results much more quickly than PCR. Instead of relying on thermocycling up to 95 °C, isothermal amplification techniques leverage unique enzymes and/or primer design for nucleic acid replication at a single temperature of 60 °C, or below. The elimination of thermocycling also removes the inherent annealing/extension synchronization that gives qPCR its precise quantification capability, making sample quantification via isothermal amplification challenging.^13^ Loop-mediated isothermal amplification (LAMP) is a popular amplification method that is performed at 60-65 °C for 30-60 minutes. Several groups have related various metrics, such as real-time reaction turbidity or end-point colorimetric indicator dyes to input nucleic acid copy numbers.^14,15^ Recombinase polymerase amplification (RPA) is a particularly attractive amplification method at the point-of-care because it is performed at a single low temperature (39 °C) and produces results in less than 20 minutes. Most RPA assays use bulk fluorescence measurement (e.g. time-to-threshold) for nucleic acid quantification, though this approach lacks precision and calibrations can vary across target genotype/subtype.^16–19^ Challenges in quantification by RPA have been attributed to desynchronized amplification, chemical initiation of the reaction, and high viscosity reaction chemistry.^13^ Some efforts to use RPA for quantification, such as those shown by Crannell *et al*. and Bender *et al*., have had some success, though there are no careful studies showing that RPA has sufficient quantitative precision for a clinical application.^20,21^

Isothermal amplification has also been used in digital amplification schemes using individual droplets or wells in microfluidic devices for absolute quantification of nucleic acids.^22–24^ Similar to ddPCR, these digital methods use binary determination of amplification in each individual well or droplet combined with Poisson distribution statistics. This approach requires that individual small aqueous reaction volumes be generated, for example, using a microfluidic device for droplet generation in a immiscible oil carrier phase,^25^ or sliding chip that compartmentalizes small volumes in wells, such as the SlipChip.^22,26^ These digital isothermal amplification methods demonstrate repeatable outputs with precise and accurate quantification, though they all require specialized chips, emulsion generators, and/or complex loading procedures that potentially increase costs and complicate POC applications.

Paper-based POC NAATs devices have leveraged isothermal amplification within commercially available porous substrates.^21,27–30^ This approach has been developed with the goal of reducing test cost and complexity as well as increasing their robustness. Lateral-flow readout has been used with some success in quantification.^31,32^ Fluorescence-based assays with various isothermal amplification chemistries have also been used in paper-based NAATs, with fluorescent signals related to sample concentration, similar to tube-based assays.^21,27,29,33,34^ In these examples, particularly in the previous works using RPA in paper membranes, fluorescent signals (through bulk fluorescent magnitude measurement and time-to-threshold) have been related to sample concentration, though with limited success.

In this work, we report a paper-based, isothermal nucleic acid amplification test that quantifies input HIV-1 RNA and DNA by leveraging distinct regions of fluorescent amplification products. We show that RPA reactions in paper membranes produce discrete amplification nucleation sites and that the number of amplification sites correlates to input NA concentrations. We develop and use image analysis algorithms to quantify RNA and DNA in the range of 67-3,000 copies per reaction in less than 20 minutes at a constant 39 °C. We demonstrate a mobile phone-based image capture system with onboard image processing using a custom Android app, showing that this method may be well suited for point-of-care applications.

## 2. Methods

### 2.1 Amplification on Membranes

RPA and RT-RPA reactions are performed on a variety of commercially available porous membranes (Millipore GF041, Whatman GF/DVA, Whatman Fusion 5, and Millipore PES GPWP04700). We first cut the membranes with a flatbed cutter (FCX4000-50ES, Graphtec America, Inc., USA) into squares to accommodate a 50 μL RPA reaction volume. Dimensions of the squares differed with respect to each membrane’s respective water absorbency. The Millipore PES GPWP04700 membrane tears when cut by the flatbed cutter, so these membranes are cut with a CO_2_ laser (PLS6.150D, Universal Laser Systems, USA). The low water absorbency of the PES membrane necessitates a square pad with a 25 μL capacity to prevent an excessively large imaging area.

Amplification pads are placed into a 60 x 15 mm polystyrene Petri dish (25384-092, VWR, USA), and a 50 μL RPA reaction (mastermix and target) is pipetted evenly onto the pad (25 μL in the case of PES). The amplification pad is then covered with PCR tape (TempPlate RT Select Optical Film, USA Scientific, USA), ensuring good sealing adhesion between the PCR tape and the Petri dish. We lid the Petri dish and seal along its edges with Parafilm M (Millipore Sigma, USA) to prevent contamination via aerosolized amplification products. The Petri dish is placed on a resistive heater (Mr. Coffee, USA) that is set to 39 °C using an external PID temperature controller with a Type-K thermocouple attached to the heating surface. The reaction is imaged for 20 minutes and then the Petri dish is sealed in a plastic bag and disposed of. The experimental process is shown in Figure 1.

**Figure 1:**
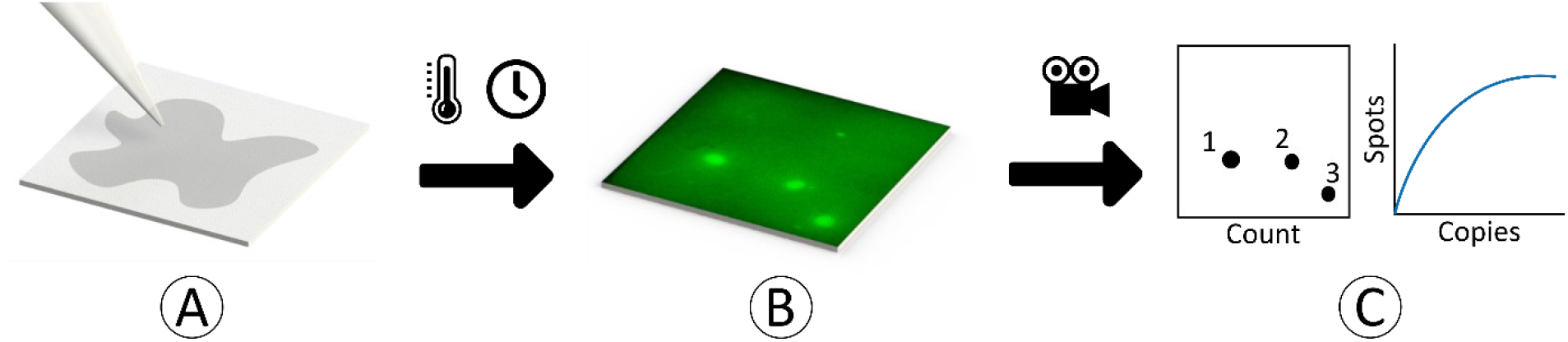
Process flow for amplification nucleation site quantification and analysis. (A) Recombinase polymerase amplification mastermix and target are dispensed onto paper membrane. The membrane is covered with PCR tape and heated to 39 °C for 20 minutes. (B) Discrete amplification nucleation sites begin to form and are recorded via fluorescent microscopy. (C) The images are analyzed by an image analysis algorithm which quantifies the number of amplification nucleation sites. This value is related to the original sample target concentration.

### 2.2 RPA and RT-RPA Conditions

In RPA experiments, we targeted synthetic DNA (gBlocks Gene Fragments, Integrated DNA Technologies, USA) that contains 1,000 base pairs of the HIV-1 genome (Group M, Subtype A). For RT-RPA experiments, we targeted purified HIV RNA from HIV-1 supernatant, prepared as described in Lillis *et al*. ^19^ Briefly, HIV-1 supernatant (Group M, Subtype A, NCBI accession number: JX140650) was received from the External Quality Assurance Program Oversight Laboratory at Duke University,^35^ and extracted using QIAamp Viral RNA Mini Kits (Qiagen, DEU). The resulting viral RNA was quantified via quantitative real-time PCR. Both target types were diluted with DEPC-treated water to create dilution series used in establishing quantifiable ranges for amplification nucleation site analysis.

In both RPA and RT-RPA, we use primers and probe developed by Lillis *et al*. for cross-subtype HIV-1 detection.^19^ The amplification mastermix consists of a TwistAmp exo kit lyophilized pellet (TwistDx, GBR), 29.5 μL rehydration buffer, 14 mM magnesium acetate, 540 nM forward and reverse primer (Integrated DNA Technologies, USA), and 120 nM exo-probe (LGC Biosearch Technologies, GBR). In the case of RT-RPA experiments, 0.08 U/μL reverse transcriptase is added (OmniScript, Qiagen, DEU). The TwistAmp exo kit instructions are otherwise followed up until incubation. Briefly, the mastermix of rehydration buffer, primers, probe, and RT (in the case of RT-RPA) is added to a lyophilized exo-kit RPA pellet, rehydrating it. We then add the target (DNA or RNA) to the mastermix and add 2.5 μL MgOAc (280 mM) to the cap lid of the tube. The tube is then closed and shaken manually for ~30 seconds to start the reaction and ensure homogenous distribution of reactants. We then immediately open the tube and pipette the 50 μL reaction volume evenly onto an amplification pad. In the case of tube-based experiments, the tubes are placed into a T16-ISO instrument (Axxin, USA), which incubates the tubes at 39 °C and records fluorescence for 20 minutes, with a manual mixing step at 4 minutes.

### 2.2 Imaging and data analysis

An epifluorescence fluorescence microscope (AZ100, Nikon, JPN) with 0.5x objective and illumination system (X-Cite exacte, Excelitas Technologies, USA) images the RPA nucleation site evolution in the pad. We use an epifluorescence filter cube set (XF100-2, Omega Optical, LLC., USA) and capture grayscale images every second for 20 minutes using a CMOS camera (Prime BSI Express, Teledyne Photometrics, USA).

A custom image analysis algorithm (MATLAB, MathWorks, USA) counts the number of amplification nucleation sites over the full 1,200 frame image-stack. The algorithm first resamples the image-stack, effectively doubling the pixel density, then averages the pixel intensity over every 10 frames to reduce image noise. It applies background subtraction to the entire image-stack by averaging the frames in the first 2 minutes and subtracting this resulting average frame from all subsequent frames on a pixel-by-pixel basis to eliminate any artifacts caused by auto-fluorescence of the amplification membranes, PCR tape, or Petri dish. A circle-finding function (based on a Circular Hough Transform algorithm) identifies discrete amplification nucleation sites for each frame. A moving-mean smoothing function averages the number of amplification nucleation sites over time using a 5-frame window, as inherent pixel noise can result in deviations in number of identified sites from frame to frame. The maximum smoothed number of sites identified over the 20 minutes is used as the number of amplification nucleation sites for that experiment. We refer to this method as the CHT (Circular Hough Transform) method. An alternative algorithm that relies on thresholding to identify the nucleation sites is also used. This algorithm (coded via ImageJ^36^) averages the pixel intensity over every 10 frames, and then performs a rolling-ball background subtraction. Thresholding of the resulting image-stack is performed using Otsu’s method, followed by a watershed transformation to separate merged nucleation sites, and then analysis via an edge-finding algorithm to identify distinct nucleation sites. We refer to this method as the TAP (Threshold Analyze Particles) method. Both codes are available in the Supplementary Information.

We also use a mobile phone to image RPA nucleation sites to demonstrate the potential use of this method in point-of-care environments. We use a Pixel 4A mobile phone (Google, USA), bandpass excitation filter (FF01-466/40-25, Semrock, USA), bandpass emission filter (FF01-550/49-25, Semrock, USA), and plano-convex macro lens (37-784, Edmund Optics, USA). The excitation and emissions bandpass filters are centered at 466 nm and 550 nm, respectively, for use with the FAM fluorophore family. The plano-convex lens is positioned such that it is directly adjacent to the mobile phone’s camera, while the excitation and emission filters are positioned underneath the mobile phone flash and camera, respectively. These are held in position by a 3D-printed fixture (S3, Ultimaker, NLD), as shown in Figure 2. The experiments were conducted in a darkroom to avoid any ambient light. Images were acquired with a fixed setting of ISO 55 and an exposure of 4 seconds.

**Figure 2:**
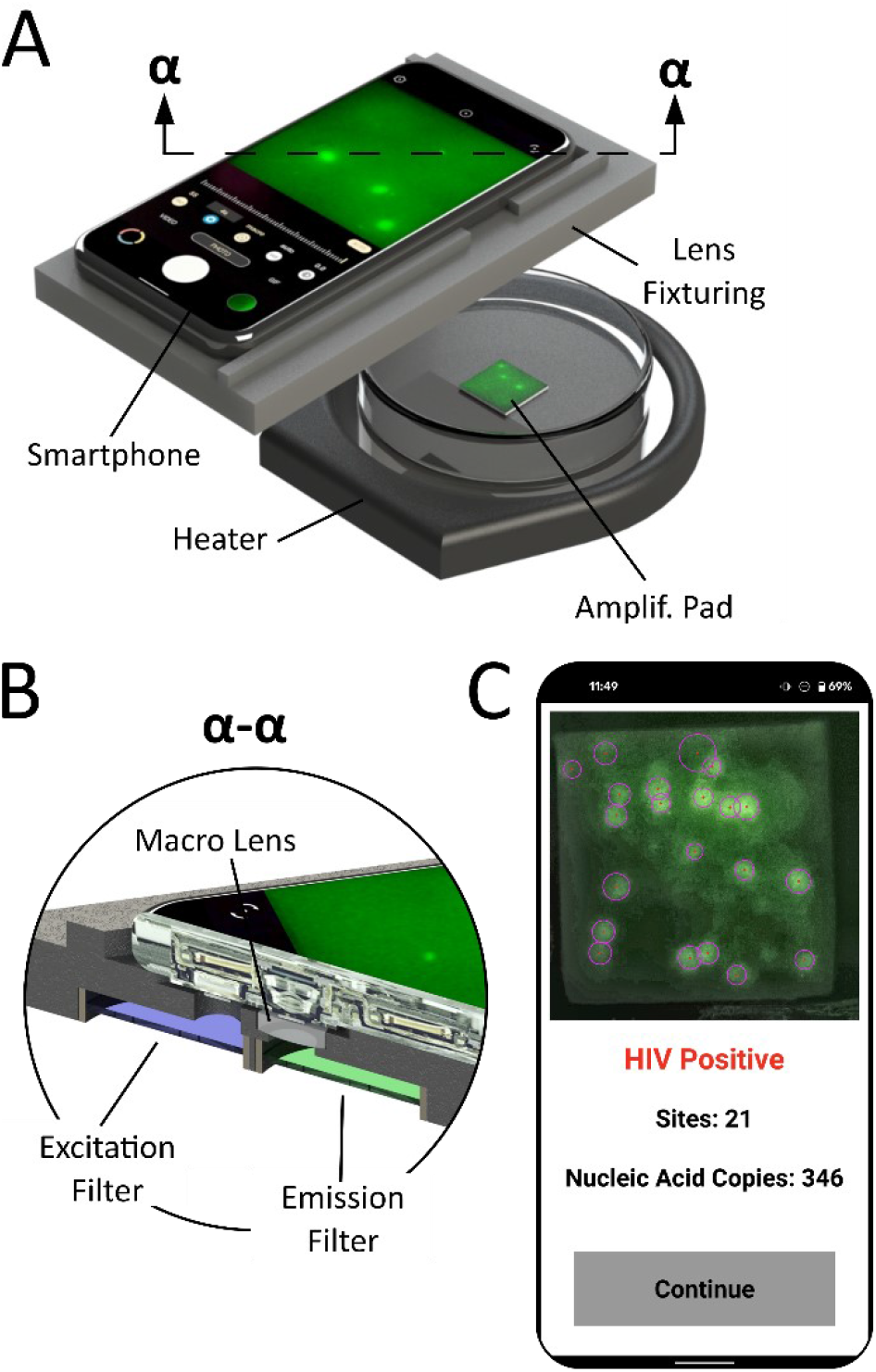
Mobile phone experimental setup for nucleation site analysis, showing (A) mobile phone, Petri dish containing the amplification membrane, heater, and 3D-printed fixturing, which holds the (B) excitation and emission filters, as well as the plano-convex macro lens. (C) Onboard processing of the images is performed using a custom app which displays the number of sites counted and calculates the estimated nucleic acid copies present.

We analyze a single frame recorded at 750 seconds into an experiment with a custom Android app written in Python 3.8.10 using the OpenCV library and run via Python for Android.^37^ The algorithm first employs an auto-cropping function to define the square region of interest containing the amplification pad. A contrast limited adaptive histogram equalization adjusts the contrast of the image, and a bilateral filter smooths the image while also removing pixel noise. The image is binarized via an adaptive Gaussian binary threshold, and a Hough circle transform then identifies and quantifies the discrete nucleation sites. The user interface displays a results screen featuring the experimental image with nucleation sites highlighted, determination of HIV status, number nucleation sites identified, and corresponding nucleic acid copies (Figure 2C).

## 3. Results and Discussion

We perform RPA and RT-RPA on GF/DVA membranes for a range of target concentrations (30-100,000 cps/rxn and 50-1,000 cps/rxn for DNA and RNA, respectively). At target concentrations of 30-3,000 cps/rxn of DNA and RNA, we observe spatially separated and distinct fluorescent amplification nucleation sites dispersed on the amplification pad that grow in diameter over the course of the experiment, as shown in Figure 3. At higher concentrations (>10,000+ cps/rxn), there are many closely packed nucleation sites that merge, resulting in splotchy heterogenous fluorescence over the amplification pad and making individual site identification more challenging.

**Figure 3:**
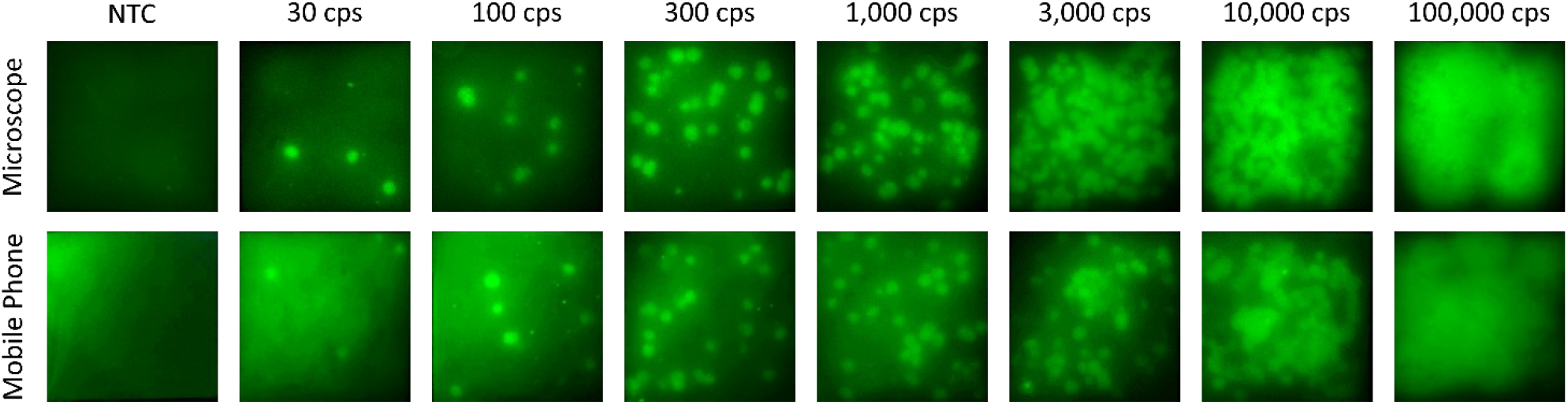
Representative fluorescence images of RPA amplification of HIV-1 DNA on GF/DVA membrane captured using a microscope and mobile phone with copy number ranging from 30 to 100,000 cps/rxn. Images were recorded at t = 750 s. At lower copy numbers (30-3,000 cps/rxn), we observe discrete amplification nucleation sites, with a positive relationship between number of nucleation sites and input copy number. At high copy numbers (10,000 and 100,000 cps/rxn), the nucleation sites are numerous and they merge into a splotchy heterogenous fluorescence image, making quantification difficult.

To our knowledge, this is the first instance that individual amplification nucleation sites in paper membranes have been described in the literature. We did not evaluate other amplification chemistries, though other publications that use these methods (e.g. LAMP, iSDA, etc. on paper membranes) exhibit homogenous fluorescence increase and not discrete amplification nucleation sites.^29,34,38–40^ We hypothesize that the nucleation sites are a result of amplification reactants (e.g. recombinase-primer filaments) or products (i.e. duplicated DNA) diffusing slowly in and out of the amplification nucleation sites. RPA’s high viscosity reaction buffer likely impedes diffusion of reactants and products, limiting the reaction to individual nucleation sites. RPA relies on viscous crowding agents such as polyethylene glycol to increase enzyme catalytic efficiency,^41^ which traditionally necessitate a mixing step halfway through tube-based RPA assay protocols.^41,42^ Porous matrices have micron scale geometries which results in viscous dampening of internal fluid flows and resulting transport of reactants and products, further contributing to a purely diffusion-limited transport and formation of discrete amplification sites.

We find significant differences between the RPA reactions in the various membranes tested. For our initial comparison tests, we use 1,000 cps HIV-1 DNA per reaction as the target. Both the GF041 and GF/DVA membranes show robust amplification, with 34-50 amplification nucleation sites visible. The Fusion 5 only results in 1-2 amplification nucleation sites and marginal increase in overall fluorescence. The PES GPWP04700 membrane supports successful amplification, in agreement with previous studies.^27,43^ We find that both the GF041 and PES membranes exhibit lower contrast between the background and amplification fluorescence in comparison to the GF/DVA membrane. Due to the robust amplification and superior contrast, all subsequent experiments were carried out using the Whatman GF/DVA membrane. Representative images of comparison RPA reactions on the various membranes are shown in Figure S1 in the Supplementary Information.

Figure 4A shows example images CHT method, including the raw image, resampled image, multi-frame averaged image, background subtraction, and identified sites. We plot the number of nucleation sites counted for the algorithms we tested compared to the number of sites counted manually. We find that CHT, TAP, and manual count methods agree well at low copy numbers (<300 cps/rxn), where there are relatively few (<50) amplification nucleation sites. At higher DNA copy numbers (1,000 - 3,000 cps/rxn), there are differences between the algorithmic counts and manual counts, though this disagreement is scattered. At higher copy number (10,000 cps/rxn), both algorithms significantly undercount the number of sites relative to the manual count. We believe this deviation is due to the algorithms being unable to robustly identify individual nucleation sites prior to the sites merging at higher input copy numbers. This is supported by plots of identified amplification nucleation sites over time of a single experiment, shown in Figure 4C. Often, the number of nucleation sites will peak halfway through an experiment (we use the maximum as the recorded value for each respective experiment) and then begin to decrease as the sites merge, revealing this weakness in the algorithms (additional nucleation sites vs. time plots are shown in the Supplementary Information). In all subsequent microscope-based experiments, we use the CHT algorithm for nucleation site quantification.

**Figure 4:**
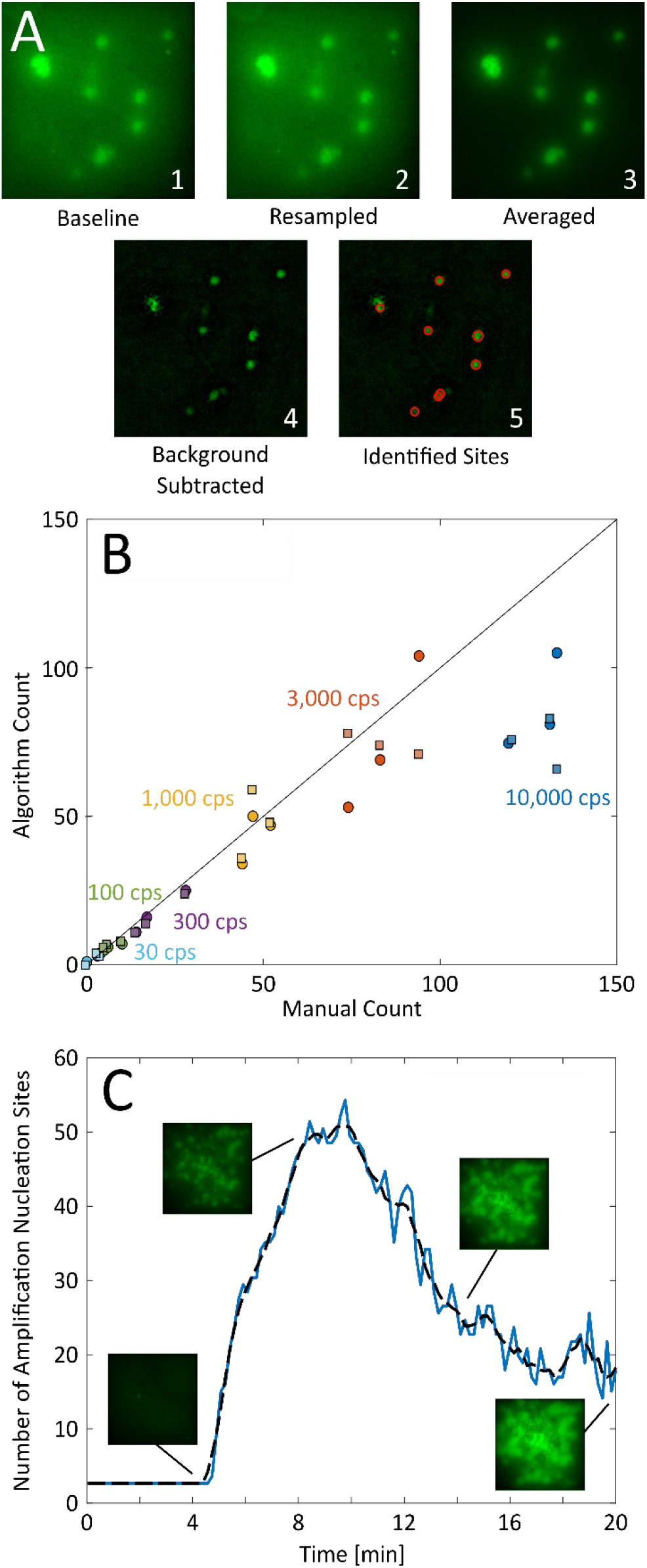
Results of algorithmic nucleation site identification and counting. (A) Circular Hough Transform (CHT) algorithm process. The raw data (1) is first resampled to increase pixel density (2) with the pixel intensity averaged over 10 frames (3). We perform background subtraction (4), subtracting the average of the first two minutes’ frames on a pixel-by-pixel basis with the resulting amplification nucleation sites then identified and quantified via a circle counting algorithm (5). (B) Comparison of the CHT (○) and TAP (□) algorithms against a manual count of RPA amplification nucleation sites with DNA target copy numbers of 30 - 10,000 cps/rxn. At lower copy numbers (≤ 3,000 cps/rxn), there is good agreement between the manual count and both algorithms. In this regime, the number of amplification nucleation sites is relatively low and there is sufficient separation between the sites for successful algorithmic quantification. At higher copy numbers (10,000 cps/rxn), the nucleation sites begin to merge, making algorithmic quantification more difficult and resulting in an undercount relative to the manual count. This undercount is observed for both algorithms, though it is more pronounced in the TAP algorithm. (C) Representative experiment of 1,000 cps/rxn DNA on Whatman GF/DVA membrane, showing the number of amplification nucleation sites as a function of experiment time, quantified via the CHT image analysis algorithm. A sharp increase in number of nucleation sites occurs near 4-10 minutes, followed by a decrease in number of identified sites as the sites begin to grow and merge. The dashed line represents the moving-mean smoothing function used.

In Figure 5 we plot the number of nucleation sites as a function of the number of nucleic acid copies per reaction for HIV-1 DNA and RNA using both the microscope and mobile phone imaging systems. The data shows that the number of amplification nucleation sites is proportional (log-log) to the number of input target copies. For RPA experiments with DNA target, input copy numbers of 30-100,000 cps/rxn were tested in triplicates and all experiments showed positive amplification except for one trial at 30 cps/rxn. No template controls (NTCs) did not show any amplification. We estimate the limit of detection (LoD) of this method via Probit analysis^44^ as copies DNA per reaction. We find that the number of amplification nucleation sites as a function of copies of DNA per reaction follows power-law relationship in the form of *y* = *b* * *x^m^*, where *b* = 0.189 and *m* = 0.762. This relationship holds well up to 3,000 cps/rxn, after which the number of measured amplification nucleation sites reaches a maximum and then decreases with increasing number of copies. The measured decrease in amplification sites is due to merging of sites at high copy numbers (10,000-100,000 cps/rxn) and the inability of the algorithms to capture and record all the nucleation sites. With our methodology, we report the dynamic range to be 67-3,000 copies of HIV-1 DNA per reaction (equivalent to 1,300-60,000 cps/mL). We show a comparison plot with a manual count in Figure S6, showing an extended dynamic range of up to 10,000 cps/rxn, suggesting that algorithm optimization can likely extend the reported dynamic range. Note that it is difficult to perform manual counts at copy numbers upwards of 10,000 cps/rxn, as the nucleation sites have already merged once they become sufficiently bright to distinguish from the background. These results suggest that alternative algorithm methodologies, such as pattern recognition or machine learning, may be useful in extending the dynamic range. While this site merging behavior limits the current dynamic range, we are still able to determine successful amplification in high copy number experiments using bulk fluorescence values.

**Figure 5:**
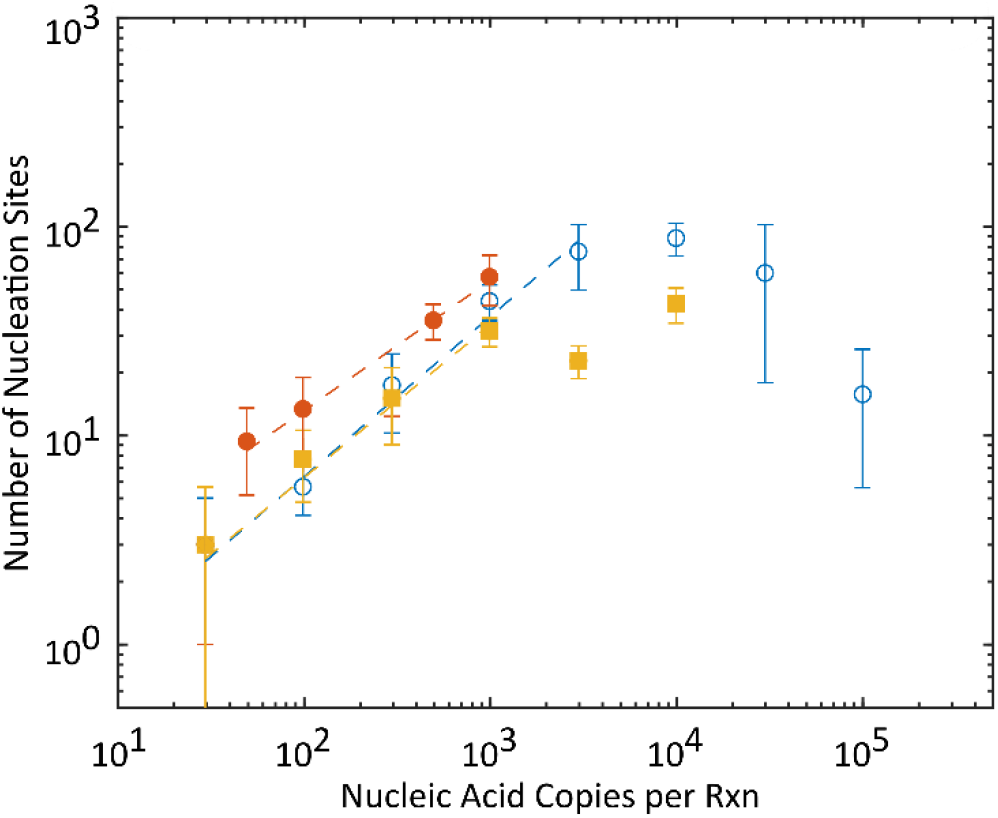
Quantification of amplification nucleation sites for HIV DNA (blue open circles) and RNA (red filled circles) on GF/DVA membrane captured using a microscope compared to HIV DNA data captured and processed using a mobile phone (yellow filled squares). Data points represent the average and standard deviations (n=3) for each target copy number tested and quantified using CHT algorithms. Dashed lines represent fitted models In all cases, we observe a strong relationship between nucleic acid input copy number and number of amplification nucleation sites. At high copy numbers (>10,000 cps/rxn), the number of measured nucleation sites decreases due to merging of amplification sites and inability of the experimental methodology to resolve and quantify the individual nucleation sites.

Similar discrete amplification nucleation sites are observed in the RT-RPA experiments with HIV-1 RNA target. The RNA copy per reaction concentration were limited to 50-1,000 cps/rxn due to the clinical specimen tested. We fit a similar power law model to the data (*y* = *b* * *x^m^*), where *b* = 0.821 and *m* = 0.590.

The mobile phone-based imaging and processing method demonstrates similar performance to microscope-based image capture for 30-1,000 cps/rxn; however, the performance of this system degrades at roughly 3,000 cps/rxn, significantly lower than the microscope-based system. This reduction in the dynamic range is due to the reduced sharpness and signal-to-noise of the mobile phone-based images and the associated challenges in counting the sites. The power law model fit to the data, ranging from 30 – 1,000 cps/rxn, is in the form *y* = *b* * *x^m^*, where *b* = 0.214 and *m* = 0.731. The fit parameters are remarkably close to the fit calculated for the microscope-acquired data.

We compare the LOD, dynamic range, time, cost, and complexity of amplification nucleation site analysis to ddPCR and various other isothermal amplification techniques in Table 1. ddPCR has the lowest limit-of-detection (1 copy per 20 μL reaction volume) and the largest dynamic range of the digital methods reviewed here (5 orders of magnitude), though requires multiple devices (droplet generator, thermocycler, and droplet reader) which cost upwards of $100,000. Other digital methods, such as the SlipChip^22^ have marginally poorer performance than ddPCR, with LODs in the range of 1,000 to 10,000 cps/mL and similar high end dynamic range. While some of these chip-based platforms may ultimately be less expensive than commercially available ddPCR systems, they are manufactured via photolithography, complicating widespread POC applications. Publications using bulk fluorescence magnitudes of isothermal amplification for quantification report dynamic ranges of 3 to 4 orders of magnitude and 20-30 minute test time. Here, we report a minimally instrumented isothermal amplification which can quantify input copies over a dynamic range of 1.5 orders of magnitude.

**Table 1.** Comparison of amplification methods to quantify pathogen concentration

| Publication | Description | Reported Range | Manufacturing | Time to Results | Quantification Scheme | Notes |
| --- | --- | --- | --- | --- | --- | --- |
| Bio-RAD QX200 | ddPCR | 5E1 to 6E6 cps/mL | | 120 minutes | Digital Poisson statistics | Requires separate droplet generator, thermocycler, and reader, \$100,000 cost |
| Shen *et al.*<sup>22</sup> | Digital RPA performed on SlipChip microfluidic chip | 1.4E3 to 1E6 cps/mL | Photolithography | 60 minutes | Digital Poisson statistics | Used 1,550 nanoliter-sized wells with automatic filling |
| Li *et al.*<sup>23</sup> | Digital RPA performed on microfluidic chip | 1.3E4 to 2.9E6 cps/mL | Photolithography | 20 minutes | Digital Poisson statistics | Used 27,000 picoliter-sized wells, rather complicated loading procedure |
| Lin *et al.*<sup>24</sup> | Digital LAMP on track-etched membrane | 1.1E4 to 1.1E8 cps/mL | LAMP mastermix added directly to polycarbonate membrane | 40 minutes | Digital Poisson statistics | Leveraged membrane pore structure to discretize amplification reactions |
| Liu *et al.*<sup>34</sup> | Paper-based LAMP | 1E6 to 1E10 cps/mL | Lyophilized reagents on glass fiber membrane | 30 minutes | Time-to-threshold analysis | Developed separate reader to measure real-time fluorescence |
| Seok *et al.*<sup>29</sup> | Paper-based LAMP | 5.9E3 to 5.9E6 cps/mL | Dried LAMP reagents on polyethersulfone membrane | 60 minutes | Bulk fluorescence magnitude | Multiplexed detection via multiple amplification pads |
| Ahn *et al.*<sup>27</sup> | Paper-based RPA | 1E2 to 1E5 cfu/mL | Dried RPA reagents on polyethersulfone membrane | 20 minutes | Bulk fluorescence magnitude | Multiplexed detection via multiple amplification pads, able to distinguish orders of magnitude input CFU concentrations |
| This work | Paper-based RPA | 1.3E3 to 6E4 cps/mL | RPA mastermix added directly to glass fiber membrane | 20 minutes | Amplification nucleation site analysis | No alterations to the membrane required, mobile phone-compatible |

Amplification nucleation site analysis may provide sufficient precision and accuracy for some point-of-care nucleic acid quantification applications. For example, in HIV-1 viral load monitoring the relevant quantification range of interest is 200-1,000,000 cps/mL (whereas the current dynamic range we show of 67-3,000 cps/rxn equates to 1,300-60,000 cps/mL).^10^ We note that in this work, we tested highly purified and simple systems and a complete POC diagnostic test may require significant sample preparation steps that impact the concentration of target in the reaction volume. Nucleation site counting may also provide greater precision and accuracy than tube-based RPA. As a preliminary estimate, we use the calculated calibration curve and determine the average predicted concentrations and associated standard deviations (n=3) for the collected data, similar to the process used by Crannell et al.^20^ For microscope-acquired DNA experiments within the quantifiable range (100-3,000 cps/rxn), we predict average concentrations within 0.09 log_10_(copies per reaction) of the correct concentration, with average standard deviations of 0.15 log_10_(copies per reaction). We perform similar calculations for tube-based RPA experiments, using the same assay, targets, and quantifiable ranges (tube-based time-to-threshold results are down in Figure S2), and are able to predict average concentrations within 0.22 log_10_(copies per reaction) with average standard deviations of 0.3 log_10_(copies per reaction). Averaged microscope-acquired RNA experiments using amplification nucleation site analysis are able to predict the correct concentration almost exactly, within 0.01 log_10_(copies per reaction), while tube-based RPA experiments are able to predict the correct concentration within 0.26 log_10_(copies per reaction). These calculations are shown in the Supplementary Information.

Amplification nucleation site analysis might also aid in quantification of RPA reactions that are slowed relative to calibration curves created with pristine and controlled/contrived target sequences. While RPA is well-known for its ability to robustly amply across genotypes and subtypes by withstanding primer mismatches,^13,45^ mismatches can have a significant effect on amplification rate and time-to-threshold.^19^ This compounds challenges in tube-based quantification if exact target sequence is not known *a priori*, for example, when multiple genotypes and subtypes are detected with the same assay. Amplification nucleation site analysis may address this, as there is no time dependency. Crude sample preparation with remnant inhibitors/confounders can also adversely affect traditional amplification quantification, though it is not yet clear if this will impact the quantification accuracy of amplification nucleation site counting. The impact of primer mismatches and sample preparation, in addition to determining the true precision and accuracy of amplification nucleation site analysis compared to tube-based RPA is the focus on future work.

Mobile phone- and inexpensive-based fluorescence readers have been used to detect and quantify isothermal amplification assays, typically using either endpoint fluorescence or time-to-threshold values.^38,46–48^ Mobile phone or inexpensive optical component based readers may be appealing for a POC setting because they potentially result in lower costs and greater accessibility. However, it is well known that differences in phone components (i.e. camera quality and/or flash spectrum) can result in differing signals,^49,50^ potentially confounding quantification if not properly calibrated. Amplification nucleation site analysis does not rely on relative fluorescence or precise pixel intensity values and may prove to be a more robust and consistent method of quantification across inexpensive optics as well as mobile phone types and models.

## 5. Summary

We present a novel method for HIV-1 DNA and RNA nucleic acid quantification using paper-based isothermal amplification and nucleation site amplification counting. We observe discrete fluorescent amplification nucleation sites when recombinase polymerase amplification is performed on porous fiber substrates and use an image analysis algorithm to count the sites. The number of amplification sites follows a power law relation to the number of input nucleic acid copies. Using DNA targets, we report a quantifiable range of 67-3,000 cps/rxn, as higher copy concentrations lead to significant amplification nucleation site merging and difficulty in site quantification using our algorithms. With HIV-1 RNA targets, we show well-defined correlation between nucleation sites and input copies between 50-1,000 cps/rxn. The amplification process takes less than 20 minutes at a single temperature with inexpensive materials and minimal user steps, and we believe that amplification nucleation site analysis could be leveraged in low-cost point-of-care nucleic acid amplification tests to provide more robust isothermal nucleic acid amplification quantification ability compared to current methods. We also present a mobile phone-based imaging system that is able to quantify amplification nucleation sites with a custom onboard processing app with similar performance to that of a microscope-based setup, suggesting that mobile phone-based amplification nucleation site analysis is a viable strategy for point-of-care applications. Future work includes extending the quantifiable range, comparing the precision and accuracy of this method to tube-based RPA, and verifying the system using clinical samples across genotypes.

We observe merging of nucleation sites at higher concentrations of target that limits the quantifiable range of our approach. There are several ways to improve the dynamic range. Improved image analysis algorithms, perhaps one using pattern recognition over time, may extend the quantifiable range. Dilution of the target, which is regularly used in ddPCR, will shift the quantifiable range to higher concentration values, but would require multiple amplification pads to effectively increase the dynamic range. Further optimization of membrane properties may also be useful in improving the limit-of-detection, as the important characteristics and properties of different membrane types in relation to their ability to sustain amplification reactions is still unclear.

We hypothesize that the RPA grouping agents, which increase the reaction fluid’s bulk viscosity, are crucial in the formation of individual nucleation sites. For this reason, we do not expect this method will be possible with other native isothermal amplification chemistries; however, it may be possible to increase the viscosity of other amplification techniques (e.g. LAMP) through grouping agents with similar results.

## 7. Acknowledgements

The work reported in this publication was supported by the National Institute of Biomedical Imaging and Bioengineering of the National Institutes of Health under Award Number R01EB022630 and by the National Center for Advancing Translational Sciences of the National Institutes of Health under Award Number TL1TR002318. Part of this work was conducting using equipment in the Biochemical Diagnostics Foundry for Translational Research supported by the M.J Murdock Charitable Trust. The content is solely the responsibility of the authors and does not necessarily represent the official views of the National Institutes of Health.

