## Supplementary Information for "Quantitative Isothermal Amplification on Paper Membranes using Amplification Nucleation Site Analysis"

Mechanical Engineering

Stevens Way, Box 352600

Seattle, WA 98195

**Table of Contents**

This Supplementary Information document describes:

**1.** Figure S1: Representative amplification reactions on various membranes.

**2.** Figure S2: Time-to-threshold analysis of tube-based RPA experiments

**3.** Figure S3: Representative normalized number of nucleation sites over the course of 20-minute experiments

**4.** Figure S4: Time at peak number of nucleation sites for HIV-1 DNA experiments

**5.** Figure S5: Representative experiment of 1,000 cps/rxn DNA on Whatman GF/DVA membrane captured by the mobile phone imaging system

**6.** Figure S6: Log-log plots of quantification of amplification nucleation sites for HIV DNA, using the CHT algorithm and a manual count

**7.** Concentration prediction comparison between amplification nucleation site analysis and tube-based time-to-threshold analysis

**8.** Circular Hough Transform (CHT) Method Algorithm – MATLAB Based

**9.** Threshold Analyze Particles (TAP) Method Algorithm – ImageJ Based

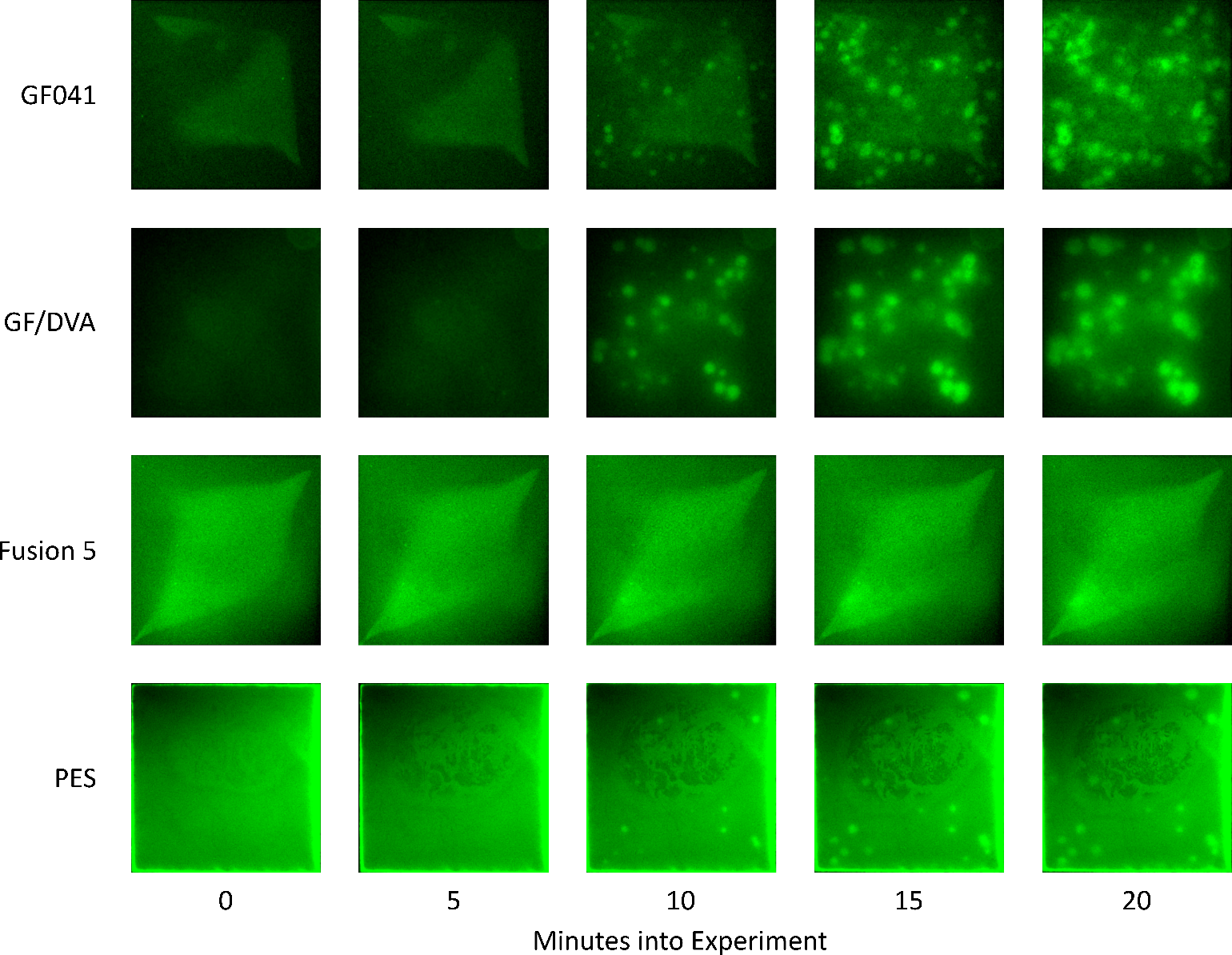

**Figure S1**: Representative amplification reactions on various membranes. From top to bottom: Millipore GF041, Whatman GF/DVA, Whatman Fusion 5, Millipore PES GPWP04700. All experiments are at 1,000 cps DNA per rxn. Due to the low water absorbency (~ 14 μL/cm^2^) of the polyethersulfone (PES) membrane, a pad to hold the full 50 μL RPA mastermix volume would be too large for the imaging set up. Consequently, only 25 μL mastermix was used for the PES experiments. Strong amplification is seen in both the Millipore GF041 and the Whatman GF/DVA, with many distinct amplification nucleation sites visible. However, the Millipore GF041 exhibits brighter background fluorescence, reducing the contrast of the amplification nucleation sites compared to the Whatman GF/DVA membrane. Amplification within the Whatman Fusion 5 is relatively poor, with only one nucleation site appearing after 20 minutes and high background signal. The Millipore PES membrane supports amplification, though air bubbles (visible in the center of the pad) would consistently form, obscuring amplification nucleation sites.

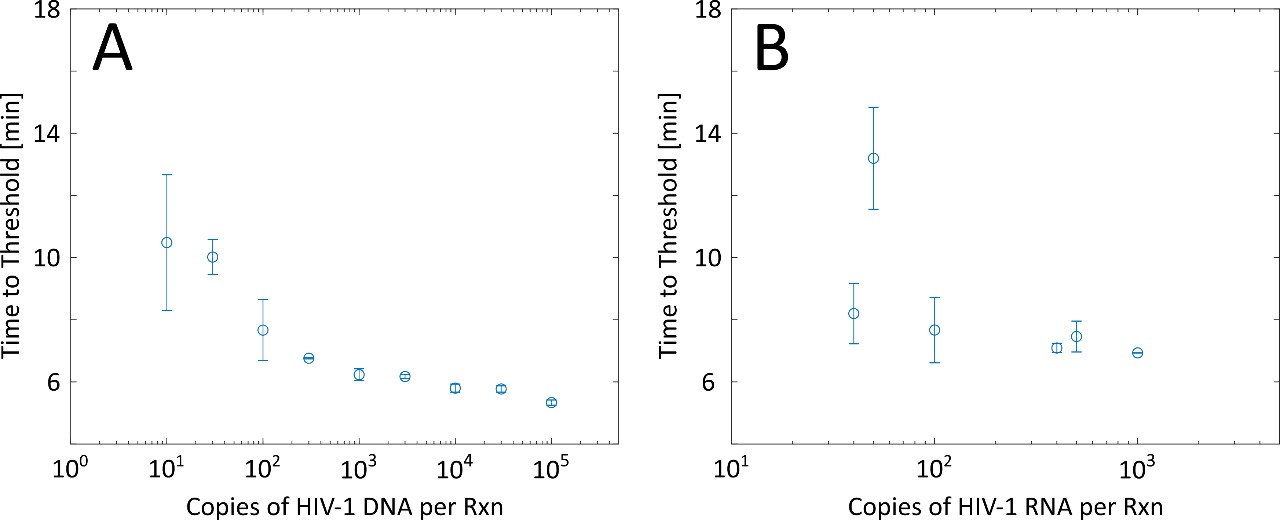

**Figure S2**: Time-to-threshold analysis of tube-based RPA experiments using the same primers and probe as used in paper-based amplification nucleation site analysis. Circles represent mean values with standard deviations shown. (A) HIV-1 DNA targets 10-100,000 cps/rxn, with each copy concentration replicated a minimum of 3 times. We observe a slight relationship between time-to-threshold and copy number, with higher copy numbers taking less time to reach the fluorescent threshold, though there is significant overlap when comparing various copy numbers, particularly at low copy numbers. (B) HIV-1 RNA targets 40-1,000 cps/rxn. We observe very little relationship between time-to-threshold and input copy number, making quantification difficult.

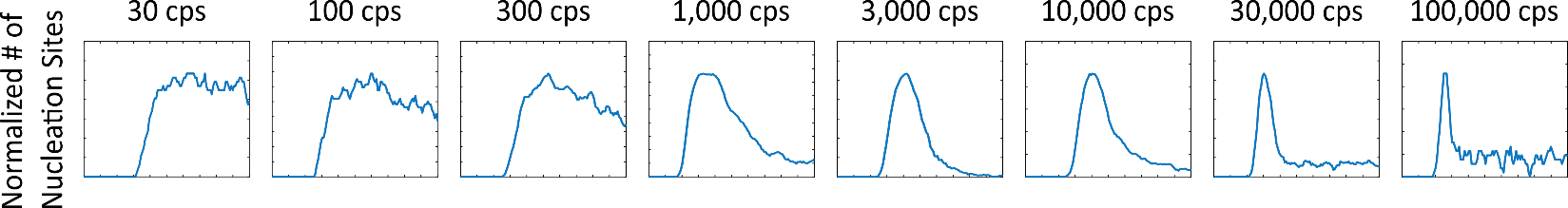

**Figure S3**: Representative normalized number of nucleation sites over the course of 20-minute experiments using HIV-1 DNA copy numbers 30 – 100,000 cps/rxn, captured via the microscope setup. Number of nucleation sites is normalized by the maximum number of nucleation sites. At very low copy numbers, the nucleation sites are spatially separated such that there is little site merging, and consequently, we do not observe any distinguishable peak of number of quantified nucleation sites. As the copy number increases, we observe gradually more pronounced peaks, as the increase in number of sites leads to more site merging. At very high copy numbers, the number of quantified nucleation sites increases rapidly, then decreases very quickly as site merging dominates very soon after the nucleation sites become visible.

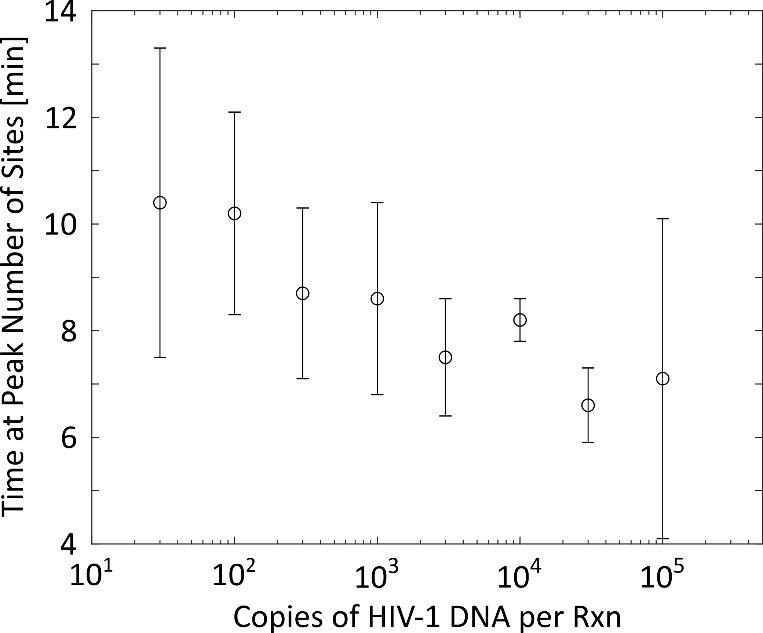

**Figure S4**: Time at peak number of nucleation sites for HIV-1 DNA experiments. As copy number increases, we generally observe the peak number of nucleation sites occur earlier in the experiment, as site merging becomes dominant very quickly at high copy numbers.

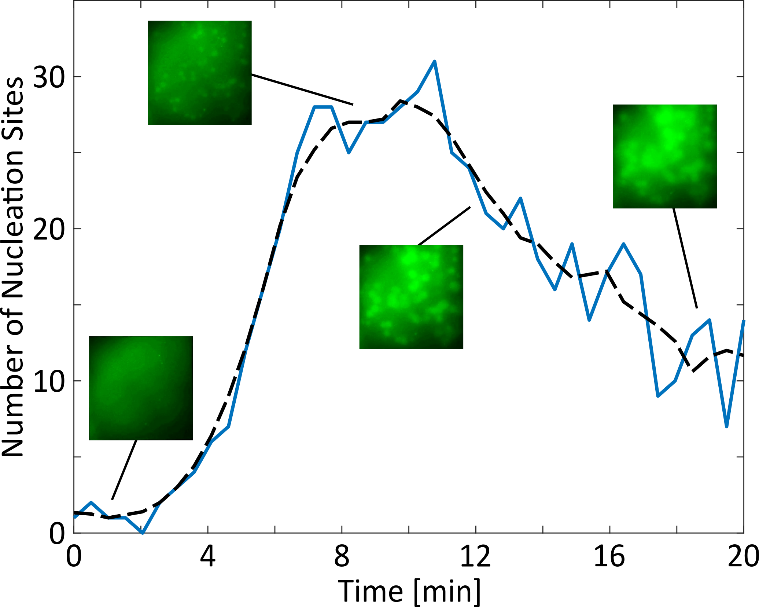

**Figure S5**: Representative experiment of 1,000 cps/rxn DNA on Whatman GF/DVA membrane captured by the mobile phone imaging system, showing the number of amplification nucleation sites as a function of experiment time. We observe very similar trends in the mobile phone-recorded experiments (in terms of site merging events) compared to microscope-recorded experiments.

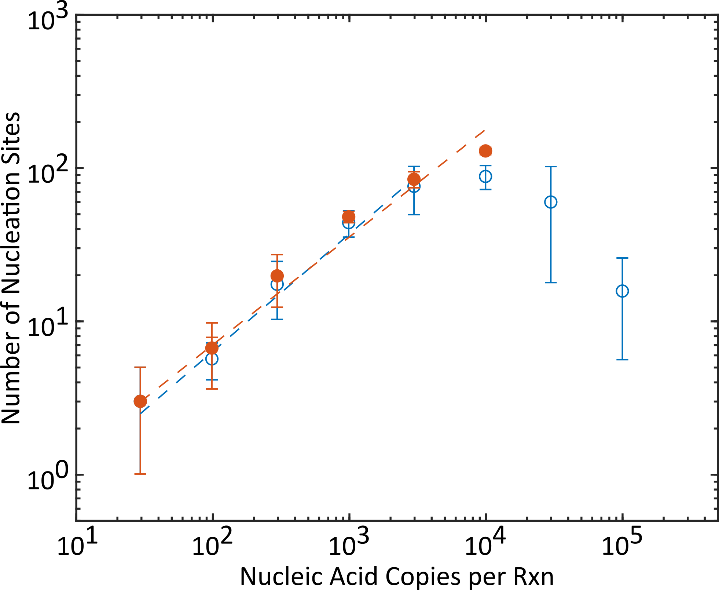

**Figure S6**: Log-log plots of quantification of amplification nucleation sites for HIV DNA, using the CHT algorithm (blue) and a manual count (red). The effects of the algorithmic undercount (relative to the manual count) are seen at 10,000 copies, suggesting that a more robust algorithm could further extend the dynamic range of amplification nucleation site analysis. Manual counts past 10,000 cps/rxn prove to be extremely difficult to perform, as there is very little separating the point in time where the nucleation sites are sufficiently bright to identify and when they begin merging.

**Concentration prediction comparison between amplification nucleation site analysis and tube-based time-to-threshold analysis**

Calibration curve fit for amplification nucleation site analysis (DNA: 100-3,000 cps/rxn):

$\# of Amplification Nucleation Sites=0.189*\left( DNA cps per rxn \right)^{0.762}$

Calibration curve fit for tube-based time-to-threshold analysis (DNA:100-3,000 cps/rxn)

$TimeToThreshold\left[ s \right]=592*\left( DNA cps per rxn \right)^{-0.062}$

Average predicted concentrations and standard deviations are calculated from triplicates (n=3). All units are in log(cps/rxn) unless otherwise noted.

| **Copies per Rxn** | **Log(Copies per Rxn)** | **Amplification Nucleation Site Analysis** | | | **Tube-Based Time-to-Threshold Analysis** | | |
| --- | --- | --- | --- | --- | --- | --- | --- |
|  |  | **Average Predicted Concentration** | **Std. Dev.** | **Error** | **Average Predicted Concentration** | **Std. Dev.** | **Error** |
| 3,000 | 3.48 | 3.39 | 0.16 | -0.09 | 3.29 | 0.07 | -0.19 |
| 1,000 | 3.00 | 3.09 | 0.10 | 0.09 | 3.22 | 0.21 | 0.22 |
| 300 | 2.48 | 2.54 | 0.19 | 0.06 | 2.65 | 0.02 | 0.17 |
| 100 | 2.00 | 1.92 | 0.13 | -0.08 | 1.82 | 0.88 | -0.18 |

Calibration curve fit for amplification nucleation site analysis (RNA: 100-1,000 cps/rxn):

$\# of Amplification Nucleation Sites=0.821*\left( cps per rxn \right)^{0.590}$

Calibration curve fit for tube-based time-to-threshold analysis (RNA:100-1,000 cps/rxn)

$TimeToThreshold\left[ s \right]=1,340*\left( cps per rxn \right)^{-0.183}$

Average predicted concentrations and standard deviations are calculated from triplicates (n=3). All units are in log(cps/rxn) unless otherwise noted.

| **Copies per Rxn** | **Log(Copies per Rxn)** | **Amplification Nucleation Site Analysis** | | | **Tube-Based Time-to-Threshold Analysis** | | |
| --- | --- | --- | --- | --- | --- | --- | --- |
|  |  | **Average Predicted Concentration** | **Std. Dev.** | **Error** | **Average Predicted Concentration** | **Std. Dev.** | **Error** |
| 1,000 | 3.00 | 2.99 | 0.06 | -0.01 | 2.78 | 0.02 | -0.22 |
| 500 | 2.70 | 2.71 | 0.06 | 0.01 | 2.71 | 0.07 | 0.01 |
| 100 | 2.00 | 2.00 | 0.3 | 0 | 2.55 | 0.26 | 0.55 |
| 50 | 1.70 | 1.70 | 0.11 | 0 | 1.26 | 0.24 | -0.44 |

**Circular Hough Transform (CHT) Method Algorithm – MATLAB Based**

```
clear all; close all; clc;

%% User Inputs
namefile = uigetfile('*tif');

%% Crop Image
figure(1)
preview=imread(namefile,1200);
imshow(preview,[0 max(max(preview))])
draw = drawrectangle('Color','b','FaceAlpha',0.01); % Select region of interest
rect = customWait(draw);
xmin = double(rect(1));
ymin = double(rect(2));
height = double(rect(4));
L = double(rect(3));

%% Compile Image Data

k = length(imfinfo(namefile)); %number of frames in data stack
for j=1:k
 Image=imread(namefile,j); % read in multiple image tiff file
 imcell=imcrop(Image,[xmin ymin L height]);

 %Resample to higher pixel density
 sz = size(imcell);
 xg = 1:sz(1);
 yg = 1:sz(2);
 F = griddedInterpolant({xg,yg},double(imcell));
 xq = (0:1/3:sz(1))';
 yq = (0:1/3:sz(2))';
 vq = F({xq,yq});

 X(:,j) = vertcat(vq(:));
end

%% Smooth Data - Average Every N Frames
[m1,n1] = size(X);
aveint = 10;
X_aveint = reshape(X, m1, aveint, n1/aveint);
X_smooth = squeeze(mean(X_aveint,2));

[m2,n2] = size(vq);
[m3,n3] = size(X_smooth);

%% Subtract Background - ATB Method

fr = 0.1*n3; %Frame that is 0.1 way through image stack
X_back = mean(X_smooth(:,1:fr),2);

X_new = bsxfun(@minus,X_smooth,X_back);
[m2,n2] = size(vq);
se = strel('disk',15);

for i = 1:k/aveint
 new = reshape(X_new(:,i),m2,n2);
 new = mat2gray(new);
 new_all(:,:,i)=new;
 background = imopen(new,se);
 newback = (new - background);

J1 = newback;
 J1_all(:,:,i) = J1;
 [centers, radii, metric] = imfindcircles(J1,...
 [8 20],'objectpolarity','bright');... % Alter for performance boost!
 radii_all{i} = radii;
 metric_all{i} = metric;
 centers_all{i} = centers;
 dots(i) = length(radii);
 % Visualize dot size increase over time
 metricbest = metric(metric>0.2);
 averad(i) = mean(radii(1:length(metricbest)));
 averad(isnan(averad))=0;


 figure(2)
 imshow(J1)

 if isempty(centers) == 0
 figure(3)
 plot(centers(:,1),-centers(:,2),'k.','MarkerSize',15)
 hold on
 end
clc
end

%% Visualize Nucleation Sites
figure(2)
imshow(new_all(:,:,k/aveint/2))
for i = 1:k/aveint
 viscircles(centers_all{i},radii_all{i},'EdgeColor','b');
end

figure(4)
imshow(new_all(:,:,k/aveint/2))
viscircles(centers_all{k/aveint/2},radii_all{k/aveint/2},'EdgeColor','b');

%% Number of Nucleation Sites Over Time
smoothdots = smoothdata(dots,'movmean',5);
round(max(smoothdots))
figure(5)
set(gcf,'Units','inches');
time = linspace(0,20,length(dots));
plot(time,dots)
hold on
plot(time,smoothdots)
ylabel('Number of Amplification Nucleation Sites')
xlabel('Time [min]')

Count = ans
```

```
%% Visualize Nucleation Sites
figure(2)
imshow(new_all(:,:,k/aveint/2))
for i = 1:k/aveint
 viscircles(centers_all{i},radii_all{i},'EdgeColor','b');
end

figure(4)
imshow(new_all(:,:,k/aveint/2))
viscircles(centers_all{k/aveint/2},radii_all{k/aveint/2},'EdgeColor','b');

%% Number of Nucleation Sites Over Time
smoothdots = smoothdata(dots,'movmean',5);
round(max(smoothdots))
figure(5)
set(gcf,'Units','inches');
time = linspace(0,20,length(dots));
plot(time,dots)
hold on
plot(time,smoothdots)
ylabel('Number of Amplification Nucleation Sites')
xlabel('Time [min]')

Count = ans
```

**Threshold Analyze Particles (TAP) Method Algorithm – ImageJ Based**

run("Grouped Z Project...", "projection=[Average Intensity] group=10");

run("Subtract Background...", "rolling=10 stack");

setSlice(48)

run("Make Binary", "method=Otsu background=Dark calculate");

run("Median...", "radius=2 stack");

run("Watershed", "stack");

run("Analyze Particles...", "size=20-Infinity exclude summarize stack");
